# Human skeletal muscle methylome after low carbohydrate energy balanced exercise

**DOI:** 10.1101/2023.01.19.524676

**Authors:** Piotr P. Gorski, Daniel C. Turner, Juma Iraki, James P. Morton, Adam P. Sharples, José L. Areta

## Abstract

We aimed to investigate the human skeletal muscle (SkM) DNA methylome after exercise in low carbohydrate (CHO) energy balance (with high fat) compared with exercise in low-CHO energy deficit (with low fat) conditions. The objective to identify novel epigenetically regulated genes and pathways associated with ‘train-low sleep-low’ paradigms. The sleep-low conditions included 9 males that cycled to deplete muscle glycogen while reaching a set energy expenditure. Post-exercise, low-CHO meals (protein-matched) completely replaced (using high-fat) or only partially replaced (low-fat) the energy expended. The following morning resting baseline biopsies were taken and the participants then undertook 75 minutes of cycling exercise, with skeletal muscle biopsies collected 30 minutes and 3.5 hours post exercise. Discovery of genome-wide DNA methylation was undertaken using Illumina EPIC arrays and targeted gene expression analysis was conducted by RT-qPCR. At baseline participants under energy balance (high fat) demonstrated a predominantly hypermethylated (60%) profile across the genome compared to energy deficit-low fat conditions. However, post exercise performed in energy balance (with high fat) elicited a more prominent hypomethylation signature 30 minutes post-exercise in gene regulatory regions important for transcription (CpG islands within promoter regions) compared with exercise in energy deficit (with low fat) conditions. Such hypomethylation was enriched within pathways related to: IL6-JAK-STAT signalling, metabolic processes, p53 / cell cycle and oxidative / fatty acid metabolism. Hypomethylation within the promoter regions of genes: HDAC2, MECR, IGF2 and c13orf16 were associated with significant increases in gene expression in the post-exercise period in energy balance compared with energy deficit. Furthermore, histone deacetylase, HDAC11 was oppositely regulated at the gene expression level compared with HDAC2, where HDAC11 was hypomethylated yet increased in energy deficit compared with energy balance conditions. Overall, we identify some novel epigenetically regulated genes associated with train-low sleep-low paradigms.

## Introduction

Performing aerobic exercise with reduced muscle glycogen via restricting dietary carbohydrate (CHO) augments the activation of the AMPK-PGC-1α signaling axis and has therefore been proposed to enhance the metabolic response and overall adaptation to exercise in skeletal muscle (SkM) tissue (reviewed in (1)). The paradigm of exercising with low-CHO availability to achieve low muscle glycogen is often referred to as ‘train-low’. To prolong the positive stimulus of low muscle glycogen without affecting daily dietary patterns, a suitable strategy is to exercise in the evening followed by low-CHO intake. In the morning, a low-CHO meal is then consumed followed by a second acute exercise session, a concept called ‘sleep-low, train-low’ (2). Within a periodized training and nutrition program, athletes usually undertake low-medium intensity exercise sessions in a low glycogen state as to maximise the exercise response and subsequent training adaptation, yet high-intensity training sessions or competition are commenced with high CHO intake and therefore high glycogen availability to ensure exercise intensity is not compromised and/or to promote optimal performance.

Despite the acute positive impact of low-CHO and reduced glycogen-induced cellular signaling in SkM following exercise, many studies have reported an acute state of low energy intake and therefore energy deficit (2-7). Indeed, continuous exercise under energy deficit can compromise exercise intensity and more chronic energy deficit can impair muscle protein synthesis (8) and is associated with negative health outcomes that may ultimately impair training adaptation (9-11). We have therefore increased dietary fat intake under sleep-low train-low conditions to achieve energy balance and prevent energy deficit whilst attempting to evoke beneficial AMPK-PGC-1α signaling (12). Despite this, achieving energy balance via increasing fat ingestion in a sleep-low train-low model did not seem to enhance the exercise-responsive molecular and metabolic markers, and seemed to impair glycaemic control the following morning compared to training in a low-CHO, energy deficit state (12). It has also been suggested that beneficial metabolic signalling following acute exercise after low-CHO feeding is blunted when increasing exogenous fat consumption in a low-CHO ‘twice-per-day’ exercise model (13).

Notwithstanding, it is still plausible that exercising under low-CHO, whilst achieving energy balance, may have an advantageous impact on alternative pathways other than canonical markers. Indeed, most of the studies to date have assessed well-characterised signalling proteins and downstream candidate gene expression levels of known markers in the metabolic response to exercise. It is currently unknown what the impact of achieving energy balance under low-CHO conditions has on the SkM response to exercise using an untargeted whole-genome ‘omi’ approach. The epigenetic modification of DNA methylation across the genome (methylome) has been demonstrated to be a dynamic response that precedes changes in gene expression in SkM after acute exercise (14-16). Both acute exercise and chronic training can predominantly decrease DNA methylation (i.e., hypomethylation) in both human and rodent SkM (14-20). This is perhaps because hypomethylation, especially in gene regulatory regions such as promoters, allows transcription factor binding to enable gene expression to occur (21, 22). Indeed, there seems to be a trend that a larger proportion of the genes that demonstrate hypomethylation are associated with a ‘gene turn on’ profile in SkM in response to resistance/strength (16, 19), high-intensity (15) and aerobic exercise (18, 23, 24). Importantly, DNA methylome studies have identified novel exercise-responsive genes in human SkM that have not been previously highlighted in mRNA/transcriptome studies (14, 25, 26). Finally, DNA methylation in SkM also seems to be sensitive to high fat dietary interventions after resistance exercise (27). However, the epigenetic response of the SkM methylome following aerobic exercise in low-CHO conditions in both energy balance and energy deficit have not been investigated. We therefore aimed to investigate the human SkM methylome after sleep-low exercise in an energy balance-high fat (EB-HF) group compared with sleep-low exercise in an energy deficit-low fat group (ED-LF) with the objective of identifying novel epigenetically regulated genes and pathways associated with train-low sleep-low paradigms.

## Methods

### Ethics

The study was approved by the Norwegian School of Sport Sciences (NIH) Ethics Committee (Application ID 01-020517) and conformed to the standards of the Declaration of Helsinki. The study was registered in the Norwegian Centre for Research Data (NSD) with reference number 54131/3/ASF. All subjects were informed about the nature of the study and possible risks involved and gave written consent prior to participating in the study.

### Participants characteristics

Nine well-trained males (tier 2/3 athletes (28)) completed the study. Participants characteristics were: VO_2max_: 66 ± 6 ml/kg/min, height: 185 ± 5 cm, body mass: 81 ± 8 kg, body fat: 17 ± 5%.

### Experimental protocol

A summary of the experimental protocol is outlined in **Figure 1**. Briefly, in a randomised, counterbalanced, crossover design, participants visited the laboratory on two separate occasions to undergo two different ‘sleep-low’ interventions which were only distinguished based on the dietary intervention. The intervention aimed at depleting SkM glycogen with cycle ergometer-based exercise in the evening of day 1 (∼18:00), followed by a low-CHO diet (to avoid muscle glycogen resynthesis). The participants slept at the same premises as the laboratory and performed a second exercise session, completed with low muscle glycogen that took place in the morning of day 2 (∼7:00). The glycogen depleting exercise elicited an energy expenditure of 30 kcal/kg fat-free mass (FFM) with alternating exercise of 2 mins at 85% of aerobic peak power output (PPO) and 2 min at 50% PPO (total duration ∼2 h). Immediately following each glycogen-depleting session, participants consumed the low-CHO meals which either completely (energy balance, high-fat; EB-HF) or partially (energy deficit, low-fat; ED-LF) replaced the energy expended during glycogen depleting exercise (see **Figure 1** and ‘dietary interventions’ section below for details on the meals). In the morning of day 2, the structured cycle ergometer exercise session that lasted 75 mins was comprised of mostly low-intensity exercise (50% PPO) but included 4 × 30 seconds and 5 × 1 minute high-intensity intervals. Further details of the experimental protocol have been published elsewhere (12).

**Figure 1.**
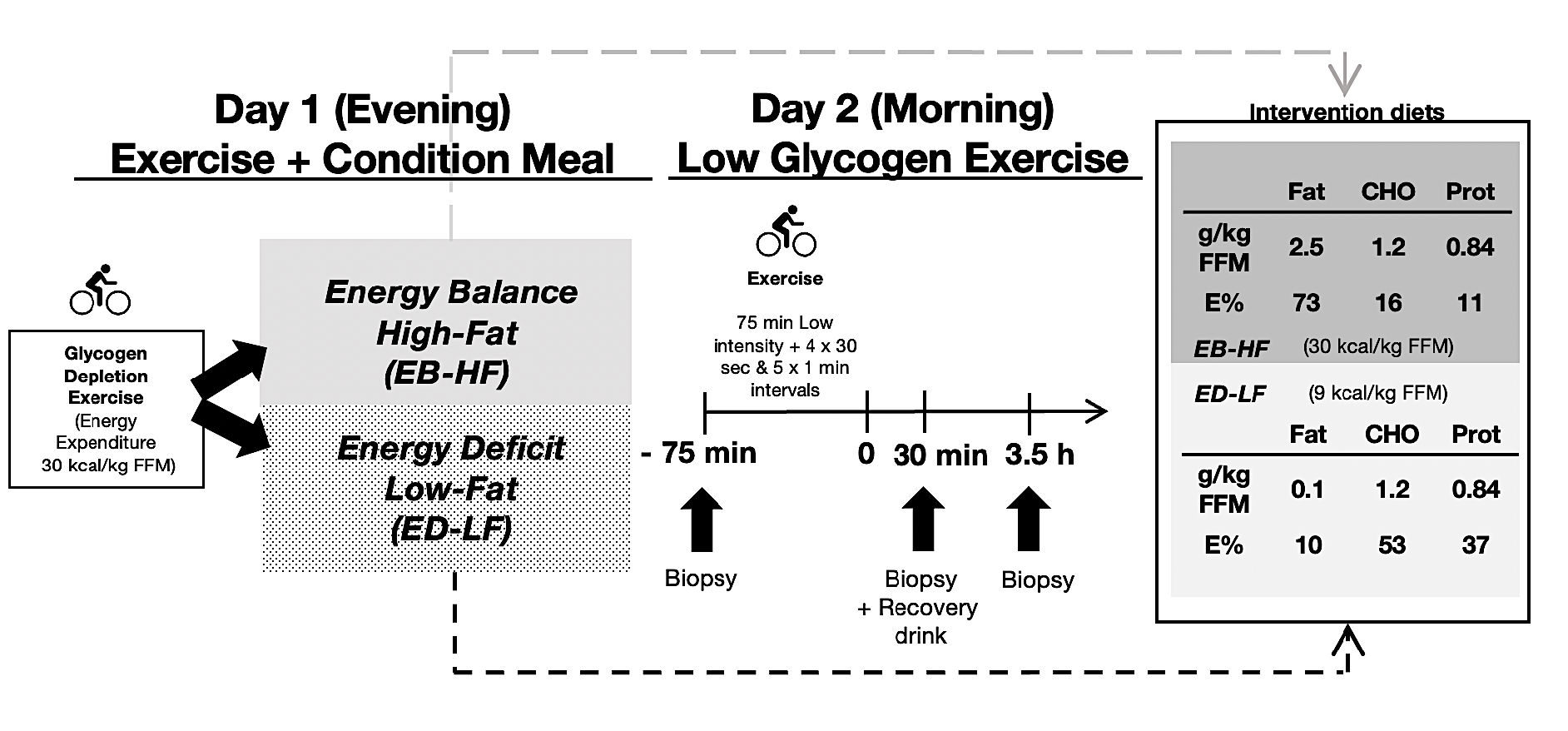
Schematic of experimental protocol and research design.

### Dietary interventions

Diet was controlled and standardised for both interventions for the 24 hrs before visiting the laboratory. Specifically, the diets were pre-packaged and provided 40 kcal/kg FFM/day containing 1.2, 6.0 and 1.35 g/kg FFM/day of fat, CHO, and protein, respectively. Immediately after the exercise in the evening of Day 1, participants consumed one of two low-CHO diets: either a high-fat, energy balance diet (EB-HF), which provided 30 kcal/kg FFM, completely replacing the energy expended during exercise and was composed of 2.5 g/kg FFM (73% energy) fat, 1.2 g/kg FFM (16% energy) CHO and 0.84 g/kg FFM (11% Energy) protein, or a low-fat, energy deficit (ED-LF) diet which provided 9 kcal/kg FFM, partially replacing the energy expended during exercise and was composed of 0.1 g/kg FFM (10% energy) fat, 1.2 g/kg FFM (53% energy) CHO and 0.84 g/kg FFM (37% energy) protein. Details of the intervention diets are also outlined in **Figure 1**. On day 2, both groups ingested a recovery drink 30 min after the morning exercise containing: 1.2 g/kg FFM CHO and 0.38 g/kg FFM of protein, as this nutrient composition is common practice for athletes to maximise training adaptation (29). Diets were designed to provide the same amount of CHO and protein while providing divergent amounts of energy (deficit and balance), with the energy difference depending solely on the difference in exogenous fat consumption.

### Biopsies

Muscle biopsies were taken from the vastus lateralis using a 6 mm Bergström needle modified for manual suction, following local anaesthesia (1% lidocaine, AstraZeneca, Cambridge, UK). Muscle biopsies were taken on day 2 at rest immediately before the start of exercise (baseline), and at 30 mins and 3.5 h after the exercise bout. From the 9 subjects completing the study, we identified a random subpopulation of 4 participants biopsies from each condition and each time point to analyse genome-wide DNA methylation (detailed methods below). Based on the genes identified to possess alterations in DNA methylation, we then validated those changes with gene expression of the same genes across the entire cohort of 9 participants (see ‘RNA isolation, primer design & gene expression analysis*’* methods section below). This helped to determine whether the identified changes at the genome-wide DNA methylation level in the subpopulation were associated with changes in gene expression of the entire cohort. Baseline characteristics of subpopulation for DNA methylome analysis were: VO_2max_: 70 ± 5 ml/kg/min, height: 185 ± 6 cm, body mass: 77 ± 6 kg, body fat: 14 ± 4%.

### Tissue homogenization and DNA isolation

Muscle samples were homogenized for 45 seconds at 6,000 rpm × 3 (5 min on ice in-between intervals) in lysis buffer (180 µl buffer ATL with 20 µl proteinase K) provided in the DNeasy spin column kit (Qiagen, UK) using a Roche Magnalyser instrument and homogenization tubes containing ceramic beads (Roche, UK). The DNA was then isolated using the DNeasy spin column kit (Qiagen, UK) bisulfite converted using the EZ DNA Methylation Kit (Zymo Research, CA, United States) as per the manufacturer’s instructions.

### DNA methylation analysis

All DNA methylation experiments were performed in accordance with Illumina manufacturer instructions for the Infinium Methylation EPIC BeadChip Array. Methods for the amplification, fragmentation, precipitation and resuspension of amplified DNA, hybridisation to EPIC beadchip, extension and staining of the bisulfite converted DNA (BCD) can be found in detail in our open access methods paper (14, 17). EPIC BeadChips were imaged using the Illumina iScan System (Illumina, United States).

### DNA methylome analysis, differentially methylated positions (DMPs), pathway enrichment analysis (KEGG and GO pathways) and differentially methylated region (DMR) analysis

Following MethylationEPIC BeadChip arrays, raw.IDAT files were processed using Partek Genomics Suite V.7 (Partek Inc. Missouri, USA) and annotated using the MethylationEPIC_v-1-0_B4 manifest file. The mean detection p-value for all samples was 0.0002, which was well below the recommended 0.01 (30). The difference between the average median methylated and average median unmethylated signal was 0.08, well below the recommended difference of less than 0.5 (30). Upon import of the data we filtered out probes located in known single nucleotide polymorphisms (SNPs) and any known cross-reactive probes using previously defined SNP and cross-reactive probe lists from EPIC BeadChip 850K validation studies (31). Although the average detection p-value for each sample across all probes was very low (on average 0.0002), we also excluded any individual probes with a detection p-value that was above 0.01 as recommended previously (30). Out of a total of 865,860 probes in the EPIC array, removal of known SNPs, cross-reactive probes, those with a detection p-value above 0.01 resulted in 809,832 probes being taken forward for downstream analysis. Following this, background normalisation was performed via functional normalisation (with noob background correction) as previously described (32). After functional normalisation, we also undertook quality control procedures via principal component analysis (PCA). One sample in the 30 min EB-HF trial was removed due to a larger variation than that expected within that condition (variation defined as values above 2.2 standard deviations for that condition). Following normalisation and quality control procedures, we undertook differentially methylated position (DMP) analysis by converting β-values to M-values (M-value = log2(β / (1 - β)), as M-values show distributions that are more statistically valid for the differential analysis of methylation levels (33). We then performed a two-way ANOVA for condition (high-fat, energy balance/EB-HF and low-fat energy deficit/ED-LF) and time (baseline, 30 minutes, 3.5 hrs) with planned contrast/pairwise comparisons of: EB-HF baseline vs. ED-LF baseline, EB-HF 30 min vs. ED-LF 30 min, EB-HF 3.5 hrs vs. ED-LF 3.5 hrs. For initial discovery of CpG sites that were deemed statistically significant, DMPs with an unadjusted P value of ≤ 0.01 were accepted for downstream analysis (Kyoto Encyclopedia of Genes and Genomes/KEGG pathway, Gene Ontology/GO and differentially methylated region/DMR analysis - see below). We then undertook CpG enrichment analysis on these DMPs within GO terms and KEGG pathways (34-36) using Partek Genomics Suite and Partek Pathway software at the significance level of FDR ≤ 0.05. Differentially methylated region (DMR) analysis was performed to identify where several CpGs were differentially methylated within a short chromosomal locations/regions, undertaken using the Bioconductor package DMRcate (DOI: <u>10.18129/B9.bioc.DMRcate</u>). Finally, to plot and visualise temporal changes in methylation across the post-exercise period (baseline, 30 min and 3.5 hr) within each condition (EB-HF and ED-LF) we implemented Self Organising Map (SOM) profiling of the change in mean methylation within each condition using Partek Genomics Suite.

### RNA isolation, primer design & gene expression analysis

Muscle tissue was homogenised in tubes containing ceramic beads (MagNA Lyser Green Beads, Roche, Germany) and 1 ml Tri-Reagent (Invitrogen, UK) for 45 seconds at 6,000 rpm × 3 (and placed on ice for 5 min at the end of each 45 second homogenization) using a Roche Magnalyser instrument (Roche, Germany). RNA was then isolated as per Invitrogen’s manufacturer’s instructions for Tri-reagent. A one-step real-time quantitative polymerase chain reaction (RT-qPCR) was performed using a QuantiFast SYBR Green RT-PCR one-step kit on a Rotorgene 3000Q thermocycler (Qiagen, UK). Each reaction was setup as follows; 4.75 μl experimental sample (7.36 ng/μl totalling 35 ng per reaction), 0.075 μl of both forward and reverse primer of the gene of interest (100 μM stock suspension), 0.1 μl of QuantiFast RT Mix (Qiagen, Manchester, UK) and 5 μl of QuantiFast SYBR Green RT-PCR Master Mix (Qiagen, Manchester, UK). Each sample was analysed in duplicate. Reverse transcription was initiated with a hold at 50°C for 10 min (cDNA synthesis) and a 5 min hold at 95°C (transcriptase inactivation and initial denaturation), before 45 × PCR cycles of; 95°C for 10 sec (denaturation) followed by 60°C for 30 sec (annealing and extension). Primer sequences for genes of interest and reference genes are in **Table 1**.

**Table 1.**
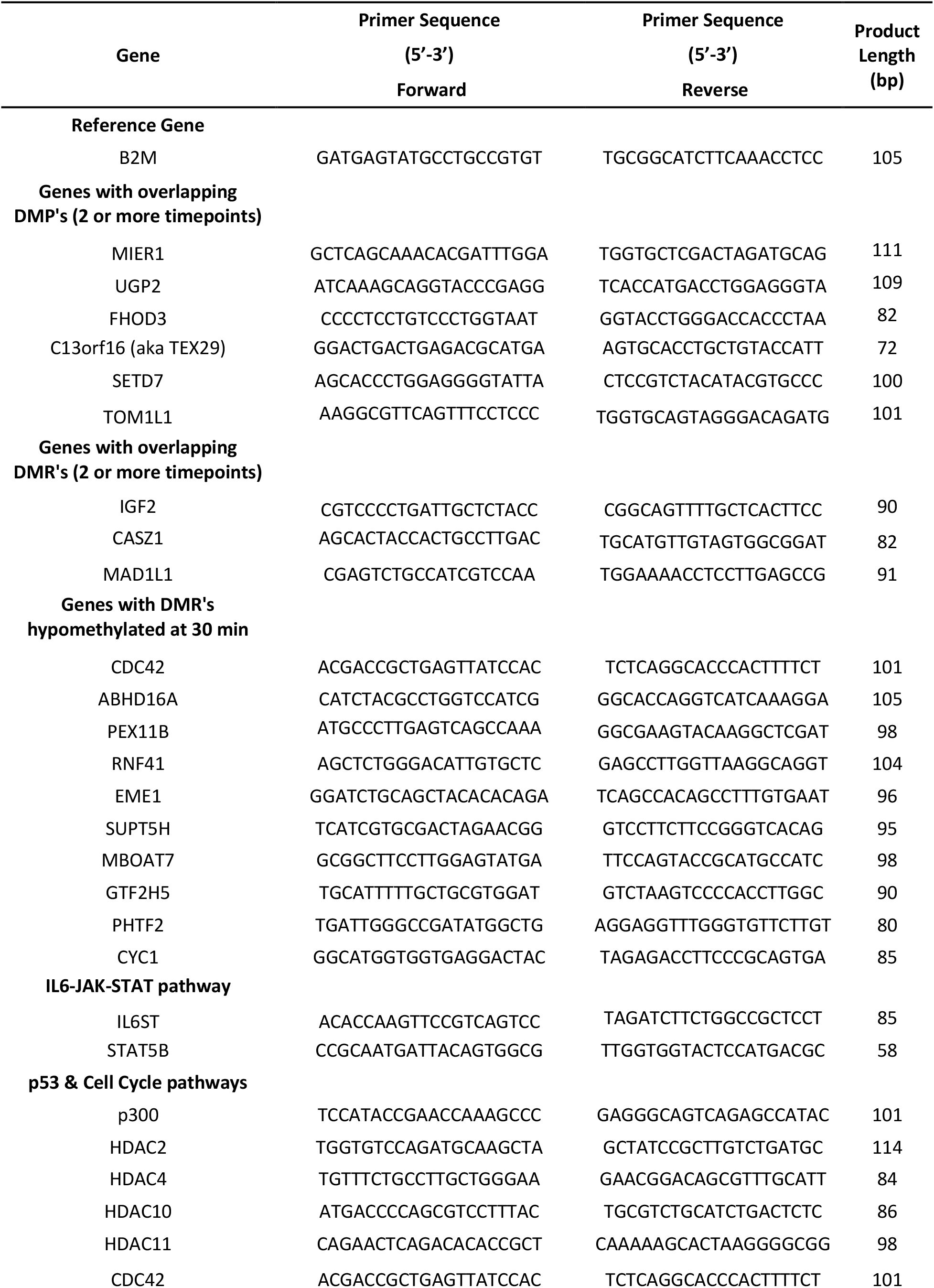
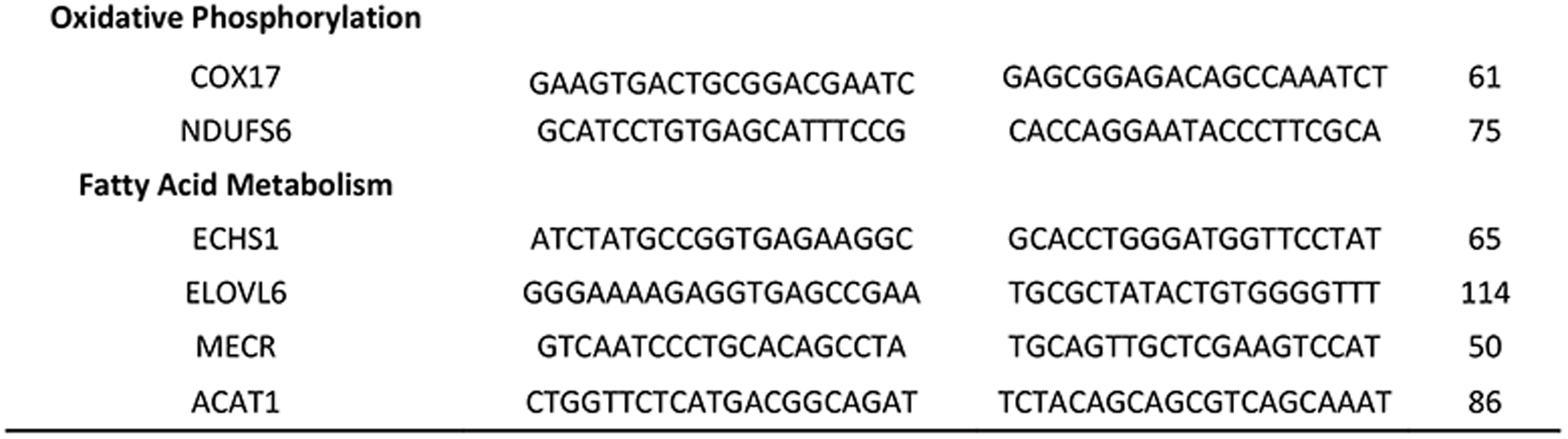
Primer sequences for RT-qPCR.

All primers were designed to yield products that included the majority of transcript variants for each gene as an impression of total changes in the gene of interest’s expression levels. All genes demonstrated no relevant unintended gene targets via BLAST search and yielded a single peak after melt curve analysis conducted after the PCR step above. All relative gene expression was quantified using the comparative Ct (^∆∆^Ct) method (37). The baseline sample for each participant was used as the calibrator condition and a pooled mean Ct was used as the reference gene (gene B2M) in the ^∆∆^Ct equation. As the average, standard deviation, and variation in Ct value for the B2M reference gene demonstrated low variation for all samples across conditions and time points (17.65 ± 0.57, 3.26% variation). The average PCR efficiencies for all the genes of interest were comparable (average of 89.76 ± 1.39%, 1.55% variation) with the reference gene B2M (88.86 ± 1.59%, 1.79% variation). Gene expression and statistical analysis was performed on n = 9 participants in both conditions and across timepoints in duplicate. Two-way ANOVA for time (baseline, 30 minutes, 3.5 hours) and condition (High fat-energy balance / EB-HF and Low fat-energy deficit/ED-LF) with Fisher LSD post hoc pairwise comparisons. Statistical analysis was performed on GraphPad Prism (version 9.2.0).

## Results

### Energy balance promotes preferential hypomethylation of gene promoter regions 30 minutes after exercise

The interaction for the 2-way ANOVA for condition (EB-HF vs. ED-LF) and time (baseline, 30 mins, 3.5 hrs) suggested there were 4,691 differentially methylation positions (DMPs) that were significantly altered across conditions and timepoints (**Suppl. File 1A**) in EB-HF relative/compared to ED-LF conditions. With main effects for condition and time identifying 4,124 and 6,662 DMPs, respectively (**Suppl. File 1B & 1C** respectively). At baseline there were 2,926 DMPs in EB-HF when compared with ED-LF **(Suppl. File 1D: Figure 2A)**, with 60% of the DMPs (1,744) hypermethylated vs. 40% (1,182) of the DMPs hypomethylated (**Figure 2B)**. At this baseline timepoint, only 11% hypermethylation and 8% of the hypomethylation occurred in CpG islands within promoters (204 out of 1,744 hypermethylated and 96 out of 1182 hypomethylated DMPs respectively; **Figure 2A and 2C; Suppl. File 1E**. At 30 minutes, in EB-HF compared with ED-LF, there was the largest total number of DMPs (9,553) identified (**Suppl. File 1F; Figure 2A**) compared with baseline (2,926 DMPs) and 3.5 hr timepoints (2,761 DMPs), with 57% DMPs hypermethylated (5,421 out of 9,553 DMPs) vs. 43% hypomethylated (4,132 out of 9,552 DMPs) (**Figure 2A and 2B)**. Importantly however, only 1% of the DMPs (51 out of 5,421) located in CpG islands within promoter regions were hypermethylated at 30 minutes, whereas 36% of all hypomethylated DMPs (1,502 out of 4,132) occurred in these important gene regulatory regions (**Figure 1A and 1C; Suppl. File 1G)**. Therefore, as a proportion of the total number of DMPs in CpG islands within promoter regions, this corresponded to 97% of the DMPs (1,502 / 1553 DMPs) possessing a hypomethylated signature at 30 minutes in EB-HF vs. versus ED-LF conditions compared with only 3% of DMPs (51 / 1553 DMPs) demonstrating a hypermethylated profile (**Figure 1D; Suppl. File 1G)**. A schematic representation of the predominant hypomethylation occurring at 30 minutes post-exercise in EB-HF is displayed as a heatmap in **Figure 2E**. Overall, this suggested that performing exercise with energy balance resulted in a preferential hypomethylation of islands and promoter regions 30 minutes post-exercise. Finally, following 3.5 hrs post exercise in EB-HF compared with ED-LF conditions, there were a lower number of total DMPs (2,761 DMPs; **Suppl. File 1H: Figure 2A**) compared with the 30-minute timepoint (9,553 DMPs), with a larger proportion of 63% DMPs hypomethylated (1,742 DMPs) versus 37% DMPS that were hypermethylated (1,019 DMPs; **Figure 2A and 2B**). Furthermore, the preferential hypomethylation of islands and promoters occurring at 30 minutes post exercise in the energy balance condition did not occur to the same extent by 3.5 hrs post exercise, where only approximately 3% (52 out of 1,742 DMPs) of the hypomethylated DMPs and 17% (171 out of 1019 DMPs) of the of the hypermethylated DMPs were in CpG islands within promoters **(Figure 2A & 1C; Suppl. File 1I)**.

**Figure 2.**
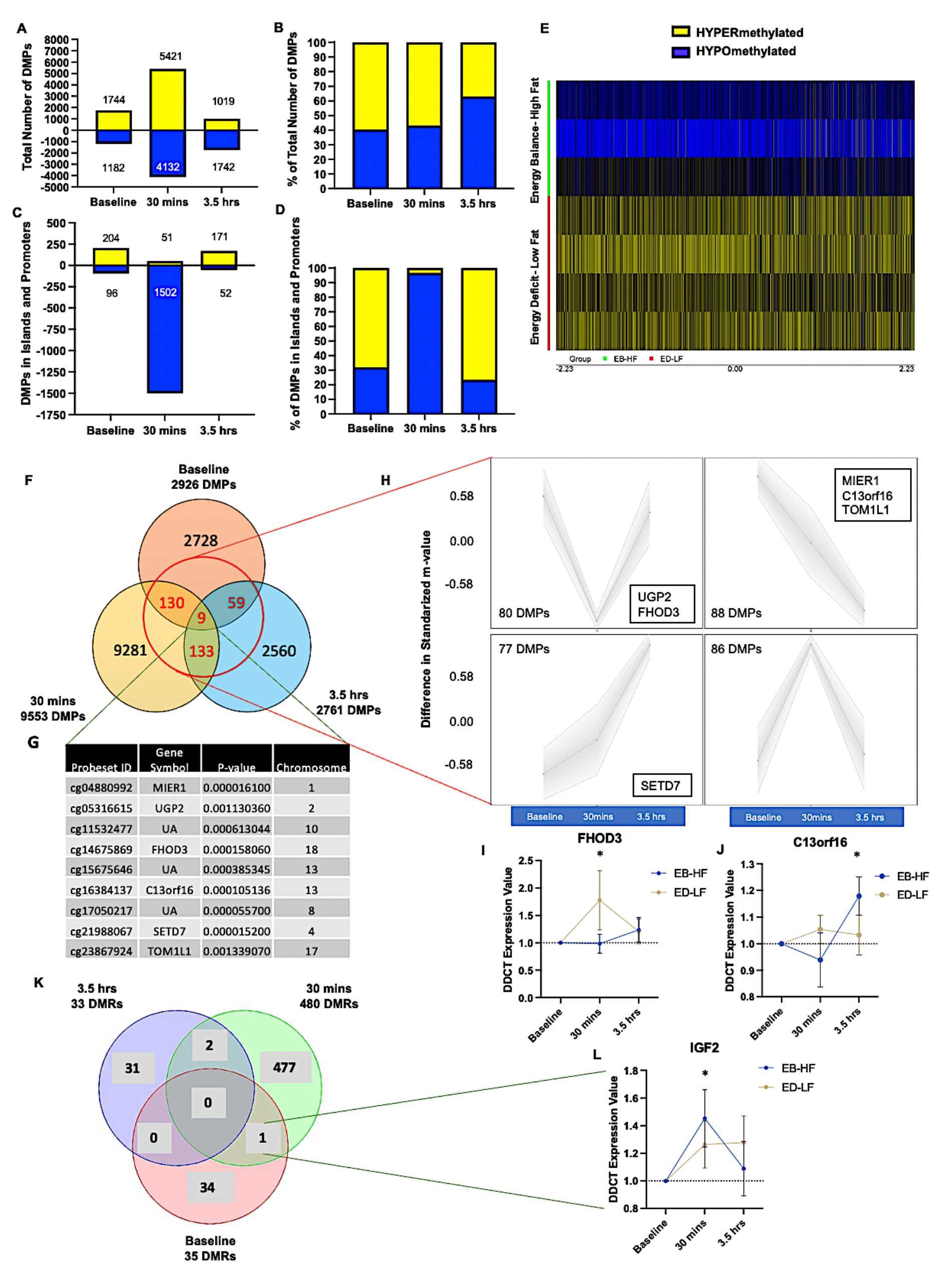
**A**. Total number of DMPs hypo (blue) and hypermethylated (yellow) between energy balance-high fat (EB-HF) compared with energy deficit-low fat (ED-LF) conditions at baseline and after 30 minutes and 3.5 hours post exercise. **B**. Hypo (blue) and hypermethylated (yellow) DMPs as a percentage of the total number of DMPs. **C**. Total number of DMPs hypo (blue) and hypermethylated (yellow) located in CpG islands within gene promoter regions between EB-HF compared with ED-LF conditions at baseline and after 30 minutes and 3.5 hours post exercise. **D**. Hypo (blue) and hypermethylated (yellow) DMPs located in CpG islands within gene promoter regions as a percentage of the total number of DMPs. **E**. A heatmap of the DMPs 30 minutes post exercise in CpG islands within promoter regions depicts that 97% (1,502 out of 1,553) DMPs demonstrated hypomethylation (blue) versus hypermethylation (yellow) in EB-HF vs. versus ED-LF conditions. **F**. Venn diagram of the 331 overlapping DMPs identified across timepoints in the EB-HF compared to the ED-LF condition. **G**. List of the 9 DMPs on 6 annotated genes altered at all timepoint comparisons in the EB-HF compared to the ED-LF conditions. **H**. SOM temporal profiling of the 331 overlapping DMPs identified across time in Figure 2F above, in the EB-HF compared to the ED-LF condition. The 6 annotated genes (from the list of the 9 DMPs in Figure 2G) have their temporal profile highlighted. **I**. Gene expression of FHOD3 identified as an overlapping DMP above. **J**. Gene expression of C13orf16 identified as an overlapping DMP above. **K**. Overlapping differentially methylated regions (DMRs) across time points in the EB-HF compared to the ED-LF condition. **L**. Gene expression of IGF2 identified as a DMR at each experimental timepoint in the EB-HF compared to the ED-LF condition in Figure 2K above. UA = unannotated to a gene.

### Gene expression of the most frequently occurring DMPs and DMRs post exercise in energy balance versus energy deficit conditions

To investigate the differentially methylated genes that were also altered at the gene expression level between conditions overtime, we identified that there were 331 DMPs that were significantly altered in the EB-HF group compared to the ED-LF group for at least 2 timepoints studied (**Figure 2F; Suppl. File 2A**). There were 9 DMPs that were altered between conditions across all timepoints, that included DMPs associated with 6 annotated genes: MIER1, UGP2, FHOD3, C13orf16, SETD7, TOM1L1 (**Figure 2G, Suppl. File 2B**). Temporal profile analysis of all 331 DMPs in the EB-HF condition that included the overlapping 9 DMPs/6 annotated genes described above, suggested that 80 DMPs, including DMPs for genes UGP2 and FHOD3, demonstrated hypomethylation at 30 minutes post-exercise that returned to baseline levels by 3.5 hrs. Furthermore, 88 DMPs including genes MIER1, C13orf16, and TOM1L1, demonstrated hypomethylation after 30 minutes and even greater hypomethylation after 3.5 hrs. Oppositely, 77 DMPs including SETD7, demonstrated a hypermethylated profile at 30 minutes and 3.5 hours in EB-HF conditions (**Figure 2H**). There were no significant differences in gene expression for genes: UGP2, MIER1, TOM1L1 and SETD7 between EB-HF and ED-LF conditions at any time point. There was however a significant increase in FHOD3 gene expression at 30 minutes post exercise (**Figure 2I**), yet this was significantly increased in the ED-LF condition and not the EB-HF condition (p = 0.05), and therefore did not inversely relate to the hypomethylated status of the gene in the EB-HF condition. There was however a significant increase (p = 0.013) in C13orf16 gene expression over time at 3.5 hrs versus 30 minutes timepoint in the EB-HF condition (**Figure 2J**), a gene that demonstrated corresponding hypomethylation at 30 minutes post exercise and even greater hypomethylation after 3.5 hrs in the EB-HF vs. ED-LF conditions. There was also an average increase of larger magnitude in the EB-HF versus the ED-LF condition at 3.5 hrs, however this did not reach statistical significance.

Gene expression is also likely to be altered if there are two or more DMPs in a short chromosomal region of a gene, known as a differentially methylated region (DMR). We therefore first undertook DMR analysis between conditions at each timepoint. There were 35, 480 and 33 DMRs identified at baseline, 30 minutes and at 3.5 hrs between the EB-HF and ED-LF conditions, respectively (**Figure 2K**; **Suppl. File 2C, D and E, respectively**). We then identified the overlapping DMRs that occurred between conditions for at least two timepoints (**Suppl. File 2F**). This included a DMR on the IGF2 gene, that was a significant DMR at baseline and 30 minutes in the EB-HF compared with ED-LF condition. As well as DMRs on genes CASZ1 and MAD1L1, identified as DMRs at both baseline and 3.5 hrs in EB-HF vs. ED-LF conditions. There were no changes in gene expression identified for CASZ1 or MAD1L1 between conditions or over time. There was, however, a significant increase in IGF2 expression in EB-HF conditions at 30 minutes post exercise compared with baseline levels (p = 0.048; **Figure 2L**), that was not significantly increased in ED-LF conditions at 30 minutes. However, this did not result reaching significance between EB-HF vs. ED-LF at the 30-minute timepoint itself.

Overall, there seems to be significantly increased gene expression in C13orf16 and IGF2 after exercise in the EB-HF condition that was not significantly changed overtime in the ED-LF condition. Albeit with the caveat that, although on average there was higher gene expression in EB-HF compared with ED-LF across these time points, this did not reach statistical significance between conditions.

### Hypomethylation at 30 minutes in energy balance conditions occurs in IL6-JAK-STAT signalling and p53 / cell cycle pathways, metabolic processes, and oxidative and fatty acid metabolism pathways

Given there were only some changes in gene expression detected for overlapping DMPs or DMRs altered across the majority of time points, we further analysed the data described above, that suggests that EB-HF leads to a preferential hypomethylation in islands within promoters compared with the ED-LF at 30 mins post-exercise. Given that gene expression changes are more likely to occur if there are more CpG sites that are differentially methylated in short chromosomal regions (especially if located in CpG islands within promoter regions), we conducted DMR analysis of CpG islands within promoters at 30 minutes in EB-HF compared with ED-LF condition. As suggested above, there were the largest number of DMRs (483 DMRs) identified at this 30-minute timepoint (**Suppl. File 2D**) with 146 DMRs within gene CpG islands and promoters (**Suppl. File 2G)**. We therefore ran gene expression of the top 10 DMRs with the most sites differentially methylated (3-4 CpG sites: **Suppl. File 2G**) in short chromosomal regions within CpG islands of promoters. This included genes: CDC42, ABHD16A, PEX11B, RNF41, EME1, SUPT5H, MBOAT7, GTF2H5, PHTF2 and CYC1 at 30 minutes in EB-HF compared with ED-LFD conditions. Where the vast majority demonstrated hypomethylated promoters at 30 minutes in EB-HF compared with ED-LF, except the gene ABHD16A that demonstrated a hypermethylated DMRs within promoters. We further identified that genes PEX11B and MBOAT7, that demonstrated hypomethylated promoter regions in EB-HF conditions, had reduced gene expression compared with ED-LF conditions (p = 0.038; **Figure 3A** and p = 0.013; **Figure 3B** respectively), where surprisingly ED-LF conditions demonstrated significantly higher gene expression at 30 minutes compared with EB-HF conditions (**Figure 3A and 3B**).

**Figure 3.**
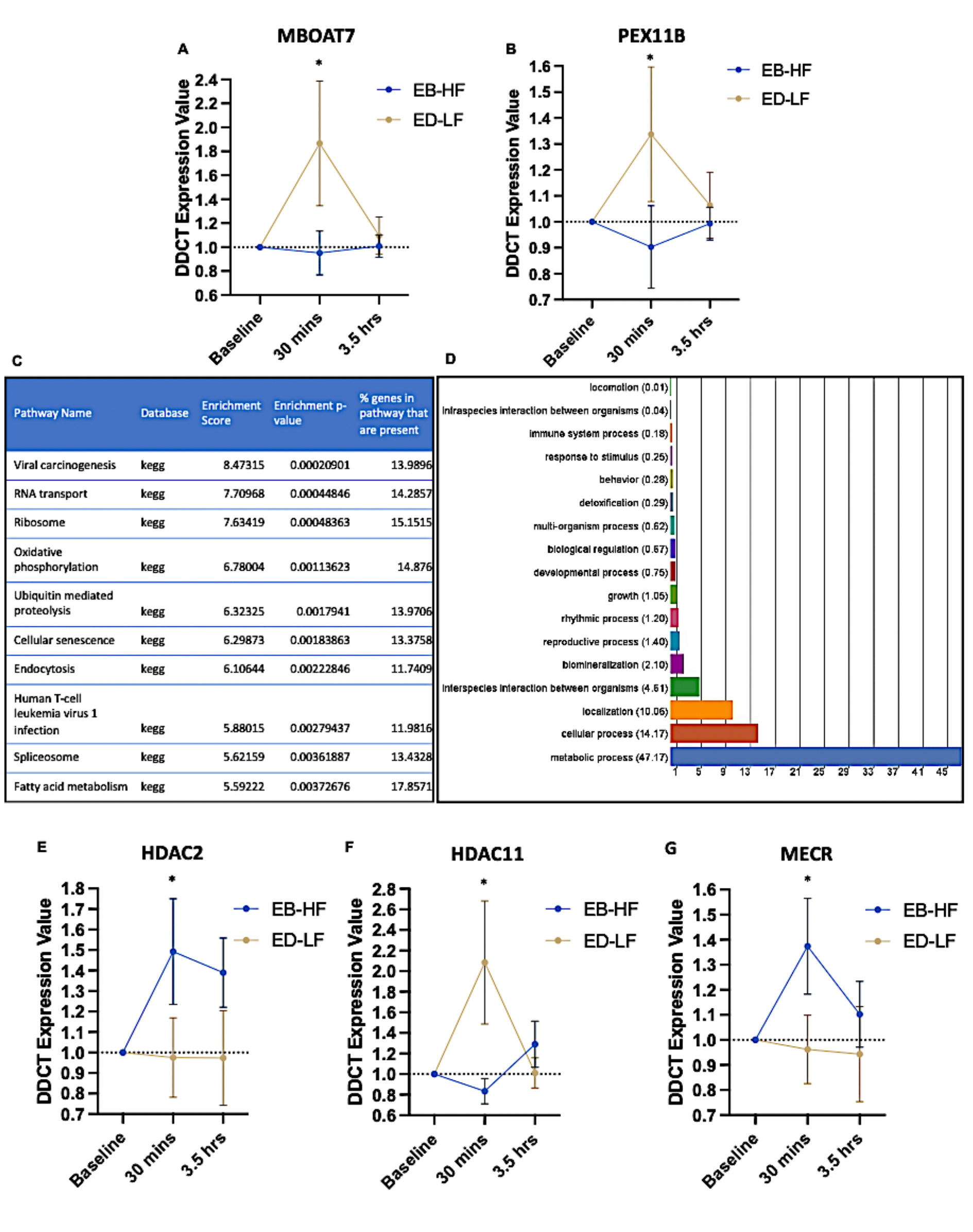
Gene expression of **A**. MBOAT2 and **B**. PEX11B identified in the top 10 most significant differentially methylated regions (DMRs) in energy balance-high fat (EB-HF) compared with energy deficit-low fat (ED-LF) conditions at 30 minutes post exercise. **C**. KEGG pathway enrichment of differential methylation at 30 minutes post exercise in CpG islands within promoter regions of genes in EB-HF compared with ED-LF conditions. With top enriched pathway ‘viral carcinogenesis’ including exercise/muscle relevant pathways IL6-JAK-STAT signalling, p53 and cell cycle pathways. **D**. GO term enrichment of differential methylation at 30 minutes post exercise in CpG islands within promoter regions of genes in EB-HF compared with ED-LF conditions. Gene expression of **E**. HDAC2 and **F**. HDAC11 identified within the top ranked KEGG pathway (Figure 3C above) and gene expression of **G**. MECR, identified within the enriched KEGG fatty acid metabolism pathway (Figure 3C above) in EB-HF and ED-LF conditions.

We therefore also undertook KEGG and GO enrichment analysis of the hypomethylated DMPs located in islands and promoters at 30 minutes. Removing non-mammalian related KEGG pathways, this identified the top 10 enriched pathways related to: Viral Carcinogenesis, RNA transport, ribosome, oxidative phosphorylation, ubiquitin mediate proteolysis, cellular senescence, endocytosis, human T-cell leukaemia virus 1 infection, spliceosome and fatty acid metabolism (**Figure 3C, Suppl. File 2H**). For GO enrichment, ‘metabolic processes’ were by far the predominant GO terms enriched for hypomethylation in the EB-HF compared with the ED-LF conditions at 30 minutes within islands and promoters (**Figure 3D; Suppl. File 2I**). We therefore first ran gene expression of genes within the most enriched hypomethylated KEGG pathway, viral carcinogenesis, as while this pathway is a disease associated pathway it includes several relevant exercise and SkM genes/pathways such as IL6-JAK-STAT signalling (for which there were hypomethylated DMPs identified on genes IL6ST, JAK1 and STAT5B), and p53 / cell cycle pathways (that included hypomethylated DMPs on genes: HDAC2, HDAC4, HDAC10 and HDAC11, CDC242 and p300). Gene expression analysis of all these genes (except JAK1 as primers demonstrated non-specificity) that all demonstrated enriched hypomethylation, we identified that HDAC2 had a significant increase in gene expression at 30 minutes post exercise (p = 0.043) in EB-HF vs. ED-LF conditions (**Figure 3E**) whereas alternatively HDAC11 significantly increased at 30 minutes post exercise (p = 0.043) in ED-LF vs. EB-HF conditions (**Figure 3F**). Given that GO term enrichment identified ‘metabolic processes’ as possessing enriched hypomethylation in islands within promoters (**Figure 3D**) we also ran gene expression for some of the most significant DMPs of those genes identified in enriched KEGG pathways related to metabolic processes (**Figure 3D**), including the oxidative phosphorylation pathway (including hypomethylated genes COX6C, COX17 and NDUFS6) and the fatty acid metabolism pathway (including hypomethylated genes ECHS1, ELOVL6, MECR, ACAT1). Gene expression for oxidative phosphorylation pathway genes COX17 and NDUFS6 were unchanged (and COX6C primers demonstrated non-specificity), as were gene expression profiles for fatty acid metabolism related genes, ECHS1 and ELOVL6. However, within the fatty acid metabolism pathway, MECR demonstrated increased gene expression at 30 minutes post exercise in EB-HF (p = 0.036) **(Figure 3G**) that was associated with the CpG island, promoter hypomethylation. Overall, achieving energy balance with high fat ingestion after exercising in low-CHO conditions preferentially hypomethylates genes in islands within promoter regions. With genes HDAC2 and MECR identified after pathway enrichment and gene expression analysis to demonstrate hypomethylated promoters and increased gene expression after exercise in energy balance compared with energy deficit conditions.

## Discussion

We aimed to investigate the genome-wide epigenetic response (via DNA methylome analysis), of human SkM after exercise in CHO restricted energy balance (via high-fat ingestion) compared with CHO restricted exercise in energy deficit (low fat ingestion) conditions. Our main objective was to identify novel epigenetically regulated genes and pathways associated with ‘train-low sleep-low’ paradigms under conditions of energy balance compared with energy deficit.

Firstly, we identified that at resting / baseline participants under energy balance (high fat) demonstrated a predominantly hypermethylated profile across the genome (60% DMPs methylated vs. 40% DMPs hypomethylated) compared to energy deficit-low fat conditions. It has been shown previously that high-fat diets can evoke hypermethylation of skeletal muscle with resistance exercise (27) and high fat ingestion for 5 days can evoke hypermethylation in human skeletal that can be maintained even when the high fat diet has ceased, and therefore over time, these epigenetic changes may lead to alterations in gene expression (38). However, the total number of differentially methylated positions at baseline was lower than the number identified post exercise at 30 minutes, and of these DMPs only a small proportion were in gene regulatory regions. Most interestingly, following 75 minutes of cycling exercise we demonstrated that increasing exogenous fat content to achieve energy balance elicited a more prominent hypomethylation of DNA in human SkM specifically 30 minutes after exercise and this occurred preferentially in gene regulatory regions (CpG islands within promoter regions) compared with exercise energy deficit with low fat consumption. After 3.5 hrs following exercise in low-CHO conditions, differential methylation across the genome was not as extensive compared with the 30 minutes post-exercise timepoint between the two dietary conditions. Previous studies have also identified that DNA methylation changes are extensive even at 30 minutes post exercise after resistance exercise (14) and also more extensive at 3 hr compared with later 6 hr timepoints (16) and also at 30 minutes compared with 24 hr time points after high intensity sprint interval exercise (15). Such early alterations in DNA methylation occur rapidly after exercise, and due to the known mechanistic role methylation has in altering accessibility and binding of transcription factors necessary for transcription, continues to supports the notion that DNA methylation precedes alterations in gene expression in the post exercise period, where gene expression typically peaks at around 3-6 *hours* post-exercise (39). The predominance of promoter hypomethylation at 30 minutes in low-CHO energy balance compared with energy deficit conditions was enriched in KEGG pathway: ‘viral carcinogensis’, that includes relevant exercise and SkM genes/pathways such as IL6-JAK-STAT signalling and p53 / cell cycle pathways. Enriched hypomethylation at 30 minutes post exercise under energy balance was also observed in important exercise regulated pathways such as; oxidative phosphorylation and fatty acid metabolism. Most importantly, we were able to identify for the first time that energy balance resulted in hypomethylation of the promoter regions of genes: HDAC2, MECR, IGF2 and c13orf16 that subsequently resulted in significant increases in gene expression in the post exercise period compared with energy deficit conditions.

Of particular interest within this study design is the gene HDAC2, where this histone deacetylase has been previously demonstrated to control metabolism and autophagy in SkM of mice, where deletion of HDAC2 can result in mitochondrial abnormalities and sarcomere degeneration (40). HDAC2 can also mediate gene expression of autophagy genes and formation of autophagosomes, such that myofibres lacking HDAC2 causes a block of autophagy and an accumulation of toxic autophagosome intermediates (40). Most relevant to the present study, mice that were fed a high fat diet from the weaning age abolished the block on skeletal muscle autophagy caused by HDAC2 deletion and prevented myopathy (40). Therefore, it may be sensible to hypothesise that HDAC2 in human SkM maybe under epigenetic control in response to higher fat ingestion in energy balance conditions and subsequently any increases in gene expression of HDAC2 would perhaps promote autophagic homeostasis. Indeed, severe energy deficit can evoke detrimental levels of autophagy (41), however, normal autophagic response after exercise is important for preserving mitochondrial function required for the recycling and disposal of macromolecules and damaged organelles (42, 43). Therefore, exercising under energy balance that is achieved via low-CHO and high-fat ingestion may function to promote autophagic homeostasis via increases in HDAC2. While this requires further investigation, what may also support this hypothesis is that genes in the endocytosis and ubiquitin mediated proteolysis pathways were also enriched for hypomethylation in islands within promoter regions at 30 minutes post-exercise in energy balance versus energy deficit conditions. Overall, perhaps suggesting that the genes in these pathways are hypomethylated under energy balance to maintain appropriate levels of recycling, degradative and disposal processes that have been demonstrated to occur under starvation, CHO restricted and energy deficit conditions. Therefore, investigating autophagy, endocytosis and ubiquitin mediated proteolysis under these conditions would be important future directions. It is also worth mentioning that another histone deacetylase, HDAC11 was alternatively regulated at the gene expression level compared with family member HDAC2, where it increased in low-CHO energy deficit compared with energy balance-high fat conditions (so was reduced in high-fat conditions relative to low fat conditions). HDAC11 is a known regulator of fatty acid metabolism and when inhibited increases oxidative fibre conversion and mitochondrial fatty acid beta-oxidation (44). Therefore, evoking energy balance with high fat after low-CHO exercise, as is the case in the present study, perhaps seems to be important in reducing HDAC11 in human SkM and may serve to promote fatty acid metabolism. However, this requires further investigation to fully confirm this mechanism.

MECR was another gene that was hypomethylated in its promoter region and increased in mRNA expression after exercise in low-CHO energy balance-high fat compared with energy deficit low-fat conditions. MECR is part of the mitochondrial fatty acid biosynthesis (mFASII) pathway and an oxidoreductase that catalyses the last step in mitochondrial fatty acid synthesis. MECR been shown to be increased at the protein level after high intensity exercise (45). However, it’s role is unknown in SkM after low-CHO exercise or in energy balance (high-fat) vs. energy deficit (low-fat) conditions. MECR has previously been linked in regulating gene expression via PPARα and PPARγ signaling and can modulate the abundance of available bioactive lipids (46, 47). It is speculative, however, given that PPAR α and γ isoforms work as fatty-acid regulated transcription factors (48), it may be that there is an interplay between the epigenetic regulation of MECR and the fatty-acid regulatory response of PPARs that warrants future investigation.

Finally, there are some limitations that warrent discussion in the present study. We also identified that there was hypomethylation within promoter regions of some genes, but surprisingly we demonstrate associated reductions in transcription of genes: FHOD3, PEX11B, MBOAT7 and HDAC11 (discussed above) in EB-HF compared with ED-LF conditions. It is also worth noting that we also analysed several more of the most significant hypomethylated DMPs and DMRs together with their corresponding gene expression level under energy balance that were not significantly altered compared with energy deficit conditions. It is plausible that the high fat ingestion may alter metabolites that are substrates for the process of methylation or transcription and may ‘break the link’ between alterations in DNA methylation leading to changes in gene transcription. Transcriptome studies would have complimented this data set to more comprehensively identify whether this trend occurs across the genome. Indeed, other studies have suggested that high fat ingestion after resistance exercise seemed to evoke considerable methylation changes without a strong overlap with alterations in gene expression (27). However, this is a speculative hypothesis and requires an assessment of metabolites known to be responsible for the process of methylation/demethylation and transcription in energy balance versus energy deficit conditions (49). Finally, the methylation data was conducted in a relatively low number of participants as a subpopulation of the entire cohort of 9 individuals. However, gene expression was conducted on the larger subpopulation to help validate the discovery of alterations in methylated gene positions/regions in the subpopulation and whether this was associated with alterations in expression of the same genes across the entire cohort.

In summary, low-CHO energy balance conditions seemed to promote an environment for enriched hypomethylation in gene regulatory regions in the post exercise period compared with energy deficit conditions. We identify some novel epigenetically regulated genes that may be involved in regulating the molecular response of skeletal muscle after train-low sleep-low exercise.

## Acknowledgments

We would like to thank the Society for Endocrinology, Norwegian School of Sport Sciences and Liverpool John Moores University who funded this project and also Jostein Hallen for his contribution to the planning and data collection.

## Data availability

DNA methylome data will be deposited and freely available via GEO **https://www.ncbi.nlm.nih.gov/geo/** once accepted in a peer reviewed journal.

## References

1. Impey SG, Hearris MA, Hammond KM, Bartlett JD, Louis J, Close GL, and Morton JP. Fuel for the Work Required: A Theoretical Framework for Carbohydrate Periodization and the Glycogen Threshold Hypothesis. Sports Med 48: 1031–1048, 2018.

2. Lane SC, Camera DM, Lassiter DG, Areta JL, Bird SR, Yeo WK, Jeacocke NA, Krook A, Zierath JR, Burke LM, and Hawley JA. Effects of sleeping with reduced carbohydrate availability on acute training responses. J Appl Physiol (1985) 119: 643–655, 2015.

3. Bartlett JD, Louhelainen J, Iqbal Z, Cochran AJ, Gibala MJ, Gregson W, Close GL, Drust B, and Morton JP. Reduced carbohydrate availability enhances exercise-induced p53 signaling in human skeletal muscle: implications for mitochondrial biogenesis. Am J Physiol Regul Integr Comp Physiol 304: R450–458, 2013.

4. Impey SG, Hammond KM, Shepherd SO, Sharples AP, Stewart C, Limb M, Smith K, Philp A, Jeromson S, Hamilton DL, Close GL, and Morton JP. Fuel for the work required: a practical approach to amalgamating train-low paradigms for endurance athletes. Physiological reports 4: 2016.

5. Morton JP, Croft L, Bartlett JD, Maclaren DP, Reilly T, Evans L, McArdle A, and Drust B. Reduced carbohydrate availability does not modulate training-induced heat shock protein adaptations but does upregulate oxidative enzyme activity in human skeletal muscle. J Appl Physiol (1985) 106: 1513–1521, 2009.

6. Yeo WK, McGee SL, Carey AL, Paton CD, Garnham AP, Hargreaves M, and Hawley JA. Acute signalling responses to intense endurance training commenced with low or normal muscle glycogen. Experimental physiology 95: 351–358, 2010.

7. Yeo WK, Paton CD, Garnham AP, Burke LM, Carey AL, and Hawley JA. Skeletal muscle adaptation and performance responses to once a day versus twice every second day endurance training regimens. J Appl Physiol (1985) 105: 1462–1470, 2008.

8. Areta JL, Burke LM, Camera DM, West DW, Crawshay S, Moore DR, Stellingwerff T, Phillips SM, Hawley JA, and Coffey VG. Reduced resting skeletal muscle protein synthesis is rescued by resistance exercise and protein ingestion following short-term energy deficit. American journal of physiology Endocrinology and metabolism 306: E989–997, 2014.

9. De Souza MJ, Nattiv A, Joy E, Misra M, Williams NI, Mallinson RJ, Gibbs JC, Olmsted M, Goolsby M, and Matheson G. 2014 Female Athlete Triad Coalition Consensus Statement on Treatment and Return to Play of the Female Athlete Triad: 1st International Conference held in San Francisco, California, May 2012 and 2nd International Conference held in Indianapolis, Indiana, May 2013. Br J Sports Med 48: 289, 2014.

10. De Souza MJ, Koltun KJ, and Williams NI. What is the evidence for a Triad-like syndrome in exercising men? Current Opinion in Physiology 10: 27–34, 2019.

11. Mountjoy M, Sundgot-Borgen J, Burke L, Ackerman KE, Blauwet C, Constantini N, Lebrun C, Lundy B, Melin A, Meyer N, Sherman R, Tenforde AS, Torstveit MK, and Budgett R. International Olympic Committee (IOC) Consensus Statement on Relative Energy Deficiency in Sport (RED-S): 2018 Update. International journal of sport nutrition and exercise metabolism 28: 316–331, 2018.

12. Areta JL, Iraki J, Owens DJ, Joanisse S, Philp A, Morton JP, and Hallén J. Achieving energy balance with a high-fat meal does not enhance skeletal muscle adaptation and impairs glycaemic response in a sleep-low training model. Experimental physiology 105: 1778–1791, 2020.

13. Hammond KM, Impey SG, Currell K, Mitchell N, Shepherd SO, Jeromson S, Hawley JA, Close GL, Hamilton LD, Sharples AP, and Morton JP. Postexercise High-Fat Feeding Suppresses p70S6K1 Activity in Human Skeletal Muscle. Medicine and science in sports and exercise 48: 2108–2117, 2016.

14. Seaborne RA, Strauss J, Cocks M, Shepherd S, O’Brien TD, van Someren KA, Bell PG, Murgatroyd C, Morton JP, Stewart CE, and Sharples AP. Human Skeletal Muscle Possesses an Epigenetic Memory of Hypertrophy. Scientific Reports (Nature) 8: 1898, 2018.

15. Maasar MF, Turner DC, Gorski PP, Seaborne RA, Strauss JA, Shepherd SO, Cocks M, Pillon NJ, Zierath JR, Hulton AT, Drust B, and Sharples AP. The Comparative Methylome and Transcriptome After Change of Direction Compared to Straight Line Running Exercise in Human Skeletal Muscle. Front Physiol 12: 619447, 2021.

16. Sexton CL, Godwin JS, McIntosh MC, Ruple BA, Osburn SC, Hollingsworth BR, Kontos NJ, Agostinelli PJ, Kavazis AN, Ziegenfuss TN, Lopez HL, Smith R, Young KC, Dwaraka VB, Frugé AD, Mobley CB, Sharples AP, and Roberts MD. Skeletal Muscle DNA Methylation and mRNA Responses to a Bout of Higher Versus Lower Load Resistance Exercise in Previously Trained Men. Cells 12: 263, 2023.

17. Seaborne RA, Strauss J, Cocks M, Shepherd S, O’Brien TD, Someren KAV, Bell PG, Murgatroyd C, Morton JP, Stewart CE, Mein CA, and Sharples AP. Methylome of human skeletal muscle after acute & chronic resistance exercise training, detraining & retraining. Scientific Data (Nature) 5: 180213, 2018.

18. Barres R, Yan J, Egan B, Treebak JT, Rasmussen M, Fritz T, Caidahl K, Krook A, O’Gorman DJ, and Zierath JR. Acute exercise remodels promoter methylation in human skeletal muscle. Cell Metab 15: 405–411, 2012.

19. Turner DC, Seaborne RA, and Sharples AP. Comparative Transcriptome and Methylome Analysis in Human Skeletal Muscle Anabolism, Hypertrophy and Epigenetic Memory. Scientific Reports (Nature) 9: 4251, 2019.

20. Wen Y, Dungan CM, Mobley CB, Valentino T, von Walden F, and Murach KA. Nucleus Type-Specific DNA Methylomics Reveals Epigenetic “Memory” of Prior Adaptation in Skeletal Muscle. Function 2: 2021.

21. Bogdanovic O, and Veenstra GJ. DNA methylation and methyl-CpG binding proteins: developmental requirements and function. Chromosoma 118: 549–565, 2009.

22. Jones PL, Veenstra GJ, Wade PA, Vermaak D, Kass SU, Landsberger N, Strouboulis J, and Wolffe AP. Methylated DNA and MeCP2 recruit histone deacetylase to repress transcription. Nature genetics 19: 187–191, 1998.

23. Rowlands DS, Page RA, Sukala WR, Giri M, Ghimbovschi SD, Hayat I, Cheema BS, Lys I, Leikis M, Sheard PW, Wakefield SJ, Breier B, Hathout Y, Brown K, Marathi R, Orkunoglu-Suer FE, Devaney JM, Leiken B, Many G, Krebs J, Hopkins WG, and Hoffman EP. Multi-omic integrated networks connect DNA methylation and miRNA with skeletal muscle plasticity to chronic exercise in Type 2 diabetic obesity. Physiological Genomics 46: 747–765, 2014.

24. Lindholm ME, Giacomello S, Werne Solnestam B, Fischer H, Huss M, Kjellqvist S, and Sundberg CJ. The Impact of Endurance Training on Human Skeletal Muscle Memory, Global Isoform Expression and Novel Transcripts. PLoS genetics 12: e1006294, 2016.

25. Seaborne RA, Hughes DC, Turner DC, Owens DJ, Baehr LM, Gorski P, Semenova EA, Borisov OV, Larin AK, Popov DV, Generozov EV, Sutherland H, Ahmetov, II, Jarvis JC, Bodine SC, and Sharples AP. UBR5 is a novel E3 ubiquitin ligase involved in skeletal muscle hypertrophy and recovery from atrophy. J Physiol 597: 3727–3749, 2019.

26. Hughes DC, Turner DC, Baehr LM, Seaborne RA, Viggars M, Jarvis JC, Gorski PP, Stewart CE, Owens DJ, Bodine SC, and Sharples AP. Knockdown of the E3 ubiquitin ligase UBR5 and its role in skeletal muscle anabolism. Am J Physiol Cell Physiol 320: C45–c56, 2021.

27. Laker RC, Garde C, Camera DM, Smiles WJ, Zierath JR, Hawley JA, and Barrès R. Transcriptomic and epigenetic responses to short-term nutrient-exercise stress in humans. Scientific reports 7: 15134, 2017.

28. McKay AKA, Stellingwerff T, Smith ES, Martin DT, Mujika I, Goosey-Tolfrey VL, Sheppard J, and Burke LM. Defining Training and Performance Caliber: A Participant Classification Framework. International journal of sports physiology and performance 17: 317–331, 2022.

29. Moore DR, Camera DM, Areta JL, and Hawley JA. Beyond muscle hypertrophy: why dietary protein is important for endurance athletes. Appl Physiol Nutr Metab 39: 987–997, 2014.

30. Maksimovic J, Phipson B, and Oshlack A. A cross-package Bioconductor workflow for analysing methylation array data [version 1; referees: 3 approved, 1 approved with reservations]. F1000Research 5: 2016.

31. Pidsley R, Zotenko E, Peters TJ, Lawrence MG, Risbridger GP, Molloy P, Van Djik S, Muhlhausler B, Stirzaker C, and Clark SJ. Critical evaluation of the Illumina MethylationEPIC BeadChip microarray for whole-genome DNA methylation profiling. Genome biology 17: 208, 2016.

32. Maksimovic J, Gordon L, and Oshlack A. SWAN: Subset-quantile within array normalization for illumina infinium HumanMethylation450 BeadChips. Genome Biol 13: R44, 2012.

33. Du P, Zhang X, Huang C-C, Jafari N, Kibbe WA, Hou L, and Lin SM. Comparison of Beta-value and M-value methods for quantifying methylation levels by microarray analysis. BMC Bioinformatics 11: 587, 2010.

34. Kanehisa M, and Goto S. KEGG: kyoto encyclopedia of genes and genomes. Nucleic acids research 28: 27–30, 2000.

35. Kanehisa M, Furumichi M, Tanabe M, Sato Y, and Morishima K. KEGG: new perspectives on genomes, pathways, diseases and drugs. Nucleic Acids Res 45: D353–d361, 2017.

36. Kanehisa M, Sato Y, Kawashima M, Furumichi M, and Tanabe M. KEGG as a reference resource for gene and protein annotation. Nucleic Acids Res 44: D457–462, 2016.

37. Schmittgen TD, and Livak KJ. Analyzing real-time PCR data by the comparative C(T) method. Nature protocols 3: 1101–1108, 2008.

38. Jacobsen SC, Brons C, Bork-Jensen J, Ribel-Madsen R, Yang B, Lara E, Hall E, Calvanese V, Nilsson E, Jorgensen SW, Mandrup S, Ling C, Fernandez AF, Fraga MF, Poulsen P, and Vaag A. Effects of short-term high-fat overfeeding on genome-wide DNA methylation in the skeletal muscle of healthy young men. Diabetologia 55: 3341–3349, 2012.

39. Egan B, and Sharples AP. Molecular Responses to Acute Exercise and Their Relevance for Adaptations in Skeletal Muscle to Exercise Training. Physiol Rev 2022.

40. Moresi V, Carrer M, Grueter CE, Rifki OF, Shelton JM, Richardson JA, Bassel-Duby R, and Olson EN. Histone deacetylases 1 and 2 regulate autophagy flux and skeletal muscle homeostasis in mice. Proc Natl Acad Sci U S A 109: 1649–1654, 2012.

41. Martin-Rincon M, Pérez-López A, Morales-Alamo D, Perez-Suarez I, de Pablos-Velasco P, Perez-Valera M, Perez-Regalado S, Martinez-Canton M, Gelabert-Rebato M, Juan-Habib JW, Holmberg HC, and Calbet JAL. Exercise Mitigates the Loss of Muscle Mass by Attenuating the Activation of Autophagy during Severe Energy Deficit. Nutrients 11: 2019.

42. Lo Verso F, Carnio S, Vainshtein A, and Sandri M. Autophagy is not required to sustain exercise and PRKAA1/AMPK activity but is important to prevent mitochondrial damage during physical activity. Autophagy 10: 1883–1894, 2014.

43. He C, Bassik MC, Moresi V, Sun K, Wei Y, Zou Z, An Z, Loh J, Fisher J, Sun Q, Korsmeyer S, Packer M, May HI, Hill JA, Virgin HW, Gilpin C, Xiao G, Bassel-Duby R, Scherer PE, and Levine B. Exercise-induced BCL2-regulated autophagy is required for muscle glucose homeostasis. Nature 481: 511–515, 2012.

44. Hurtado E, Núñez-Álvarez Y, Muñoz M, Gutiérrez-Caballero C, Casas J, Pendás AM, Peinado MA, and Suelves M. HDAC11 is a novel regulator of fatty acid oxidative metabolism in skeletal muscle. The FEBS journal 288: 902–919, 2021.

45. Granata C, Caruana NJ, Botella J, Jamnick NA, Huynh K, Kuang J, Janssen HA, Reljic B, Mellett NA, Laskowski A, Stait TL, Frazier AE, Coughlan MT, Meikle PJ, Thorburn DR, Stroud DA, and Bishop DJ. High-intensity training induces non-stoichiometric changes in the mitochondrial proteome of human skeletal muscle without reorganisation of respiratory chain content. Nature Communications 12: 7056, 2021.

46. Clay HB, Parl AK, Mitchell SL, Singh L, Bell LN, and Murdock DG. Altering the Mitochondrial Fatty Acid Synthesis (mtFASII) Pathway Modulates Cellular Metabolic States and Bioactive Lipid Profiles as Revealed by Metabolomic Profiling. PLoS ONE 11: e0151171, 2016.

47. Parl A, Mitchell SL, Clay HB, Reiss S, Li Z, and Murdock DG. The mitochondrial fatty acid synthesis (mtFASII) pathway is capable of mediating nuclear-mitochondrial cross talk through the PPAR system of transcriptional activation. Biochem Biophys Res Commun 441: 418–424, 2013.

48. Manickam R, and Wahli W. Roles of Peroxisome Proliferator-Activated Receptor β/δ in skeletal muscle physiology. Biochimie 136: 42–48, 2017.

49. Seaborne RA, and Sharples AP. The Interplay Between Exercise Metabolism, Epigenetics, and Skeletal Muscle Remodeling. Exerc Sport Sci Rev 48: 188–200, 2020.

